# Accurate background reduction in adaptive optical 3D-STED nanoscopy by dynamic phase switching

**DOI:** 10.1101/2022.06.25.497623

**Authors:** Shijie Tu, Xin Liu, Difu Yuan, Wenli Tao, Yubing Han, Yan Shi, Yanghui Li, Cuifang Kuang, Xu Liu, Yufeng Yao, Yesheng Xu, Xiang Hao

**Author notes:** **Corresponding Authors**, **Xiang Hao** – State Key Laboratory of Modern Optical Instrumentation, College of Optical Science and Engineering, Zhejiang University, Hangzhou 310027, China; Jiaxing Key Laboratory of Photonic Sensing & Intelligent Imaging, Jiaxing 314000, China; Intelligent Optics & Photonics Research Center, Jiaxing Research Institute Zhejiang University, Jiaxing 314000, China.

## Abstract

Stimulated emission depletion (STED) fluorescence nanoscopy allows the three-dimensional (3D) visualization of nanoscale subcellular structures, providing unique insights into their spatial organization. However, 3D-STED imaging and quantification of dense features are obstructed by the low signal-to-background ratio (SBR), resulting from optical aberrations and out-of-focus background. Here, combining with adaptive optics, we present an easy-to-implement and flexible method to improve SBR by dynamic phase switching. By switching to a counterclockwise vortex phase mask and a top-hat one with an incorrect inner radius, the depletion pattern features a nonzero-intensity center, enabling accurate background recordings. When the recorded background is subtracted from the aberration-corrected 3D-STED image, the SBR in dense sample areas can be improved by a factor of 3–6 times. We demonstrate our method on various dense subcellular structures, showing more advantages than the software-based background subtraction algorithms.

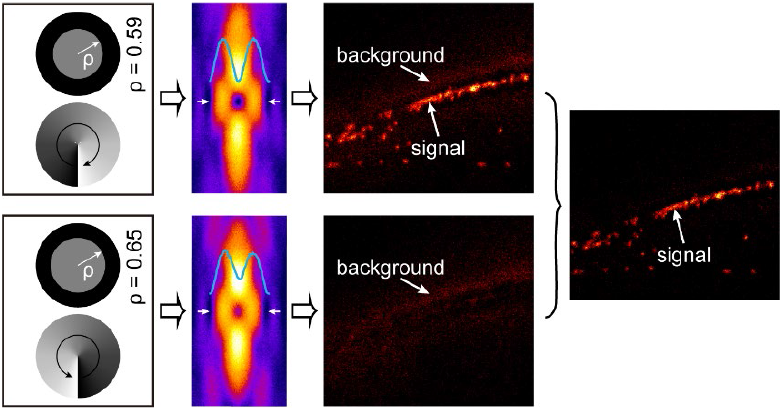

---

Over the past two decades, nanoscopy^1–3^ (or optical super-resolution microscopy) has played a key role in the noninvasive visualization of nanoscale subcellular structures and dynamics below the optical diffraction limit. Among the super-resolution imaging modalities, stimulated emission depletion (STED) nanoscopy is widely used for three-dimensional (3D) imaging of whole cells and tissues^4–6^.

To improve the axial resolution, 3D-STED nanoscopy often applies a radially symmetric “top-hat” phase mask to the depletion beam^7^, forming a point spread function (PSF) with two high-intensity lobes above and below the focal plane to deplete the axial fluorescence emission. Besides, a vortex phase mask^8^ is combined with the top-hat one to improve the lateral resolution^9^. These phase masks can be realized either by static phase plates^9^ or, more recently, by a spatial light modulator (SLM)^10^.

Compared to confocal microscopy, 3D-STED nanoscopy features a ~126-fold smaller effective focal volume^11^, but with the same detection volume. As a consequence, the fluorescence must be retained and depleted very effectively in the central and peripheral areas of the detection volume, respectively, otherwise causing a low signal-to-background ratio (SBR). However, this is not entirely achievable because the depletion PSF is sensitive to optical aberrations and has low intensity far away from the focal plane. In particular, axially extended sample areas suffer more from this background induced by the out-of-focus incomplete depletion, compared with axially thin sample areas^12^. Worse still, along with increasing the depletion power for higher resolution, the anti-Stokes excitation by the depletion laser is another background source^13^.

There have been several hardware-based or software-based techniques addressing this background issue by directly subtracting the estimated background from the regular STED images^14^. In a method called stimulated emission double depletion (STEDD), an extra Gaussian pulse is added to an ordinary 3D-STED setup to measure the background^15^. Nonetheless, STEDD is optically complex due to its requirement of an extra beam path and careful synchronization design. Polarization switching STED (psSTED), records the background by switching the polarization of the depletion beam^16^. However, psSTED is at the risk of beam path instability, using a time-consuming motorized rotation stage to switch the polarization. Worse still, it is limited in axial resolution since the higher power of STED*_xy_* beam is required to ensure accurate background recording. Another technique, applied in isoSTED nanoscopy, records the background using several offset detectors arranged symmetrically around the main detector^12^. However, this technique is hardware-complex and expensive. Moreover, by replacing the top-hat phase mask with a bivortex phase mask on the SLM, the final depletion PSF produces stronger depletion in the defocused region and therefore reduces the out-of-focus background, but with limited axial resolution^17, 18^. Besides the hardware-based techniques mentioned above, several software-based algorithms are also available for background subtraction, such as rolling ball algorithm (RBA)^19^ and wavelet-based background subtraction algorithm (WBS)^20^. However, these algorithms strongly depend on the accuracy of background estimation. Incorrect estimation potentially raises the risk of over-subtraction.

Here, we first present an adaptive optics (AO) 3D-STED nanoscopy to improve the signal via aberrations correction. Then we introduce dynamic phase switching STED (DPS-STED), a method that can accurately measure and remove the background in a 3D-STED nanoscope. The SBR can thus be effectively improved with the combination of the above two strategies.

A simplified schematic of our DPS-STED setup featuring a 100×, 1.35 numerical aperture, silicone-oil objective lens, is shown in Figure 1a (see details in Supporting Information and Figure S1). For multicolor imaging, two pulsed excitation beams and one depletion beam are used at 590-, 647-, and 775-nm wavelengths, respectively, with a 78-MHz repetition rate. For the depletion beam, we apply two orthogonal polarization components with a pulse delay from the same laser to generate lateral (STED*_xy_*) and axial (STED*z*) depletion PSFs at the sample plane without interfering with each other. A double-pass SLM configuration^21^ is used to encode the vortex or top-hat phase on each polarization component. For the excitation beams, the light from a supercontinuum laser is merged with the depletion beam via a dichroic mirror. The depletion and excitation beams then scan across the sample using a 16 kHz resonant mirror combined with two synchronized galvanometer mirrors^5^. For the detection path, fluorescence is collected by the same objective lens, descanned, and directed to a single-photon counting avalanche photodiode (APD). The measured fluorescence signals from the APDs are relayed to circuit boards for gated detection^22^. Meanwhile, a deformable mirror (DM) is placed into the detection path to correct the sample-induced aberrations.

**Figure 1.**
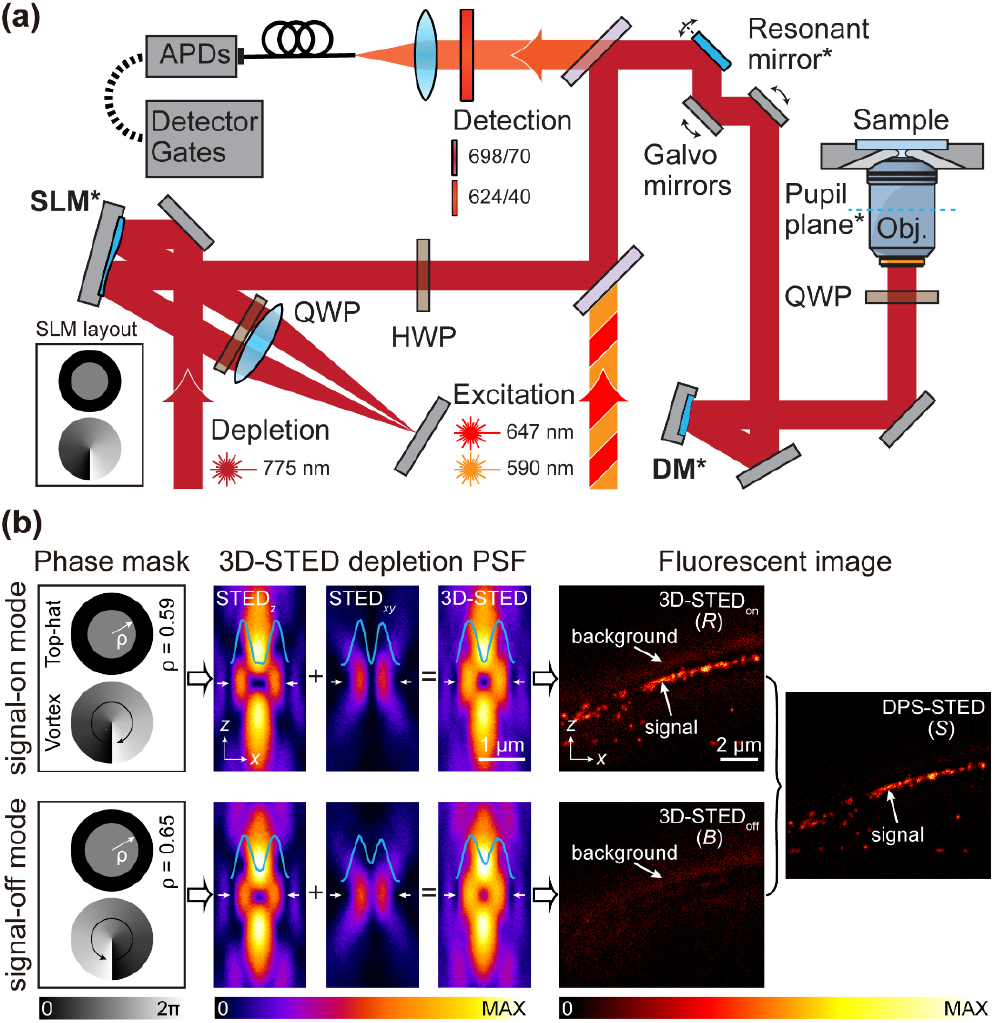
Principle of DPS-STED. (a) Simplified schematic of the optical setup. The conjugated planes are indicated with an asterisk and highlighted in blue. A detailed schematic is available in Figure S1. (b) Working principle of our method. From left to right, the sub-figures show the phase masks (pupil functions) and their effects on the depletion PSFs and the fluorescent images. The top and bottom rows are for signal-on and signal-off modes nanoscopy, respectively. The final DPS-STED result is presented on the right.

The multistep AO correction routine is shown in Figure S2. After correcting aberrations in the optical system, the shape of DM is further adjusted to compensate for the sample-induced aberrations. During the imaging, the DM is further tuned to automatically compensate for the depth-dependent aberrations^23^, an essence for imaging thick samples. In this way, our AO strategy enables aberration-free excitation and depletion PSFs at the sample, thereby enhancing the SBR via signal enhancement in confocal, 2D-STED, and 3D-STED modes. However, the background noise still hinders further image quality improvement (Figure S3).

To address this problem, we developed DPS-STED method. Its working principle is depicted in Figure 1b. In brief, DPS-STED takes effects by modulating the pupil function of the depletion beam. The STED_*xy*_ PSF generated by the vortex phase mask is polarization-dependent^24^, and the STED*_z_* PSF generated by the top-hat phase mask is determined by its inner radius^25^. Assume that the original STED_*xy*_ beam is right-handed circularly polarized. Only when the applied vortex phase mask is clockwise, a doughnut PSF with an ideally zero-intensity center is achievable^24^. In contrast, a counterclockwise vortex phase mask breaks the central “zero” by providing a strong solid longitudinal component of the electric field. Similarly, for the STED_*z*_ beam, when the top-hat phase mask only with the optimal inner radius *ρ* (this radius is determined by the intensity profile of the incident beam. In our case, *ρ* = 0.59) is applied, the STED*_z_* PSF has a zero-intensity center. Tuning the inner radius (e.g., *ρ* = 0.65) can dramatically increase the central intensity. Combining the affected STED_*xy*_ and STED*_z_* PSFs, the 3D-STED depletion PSF with almost the same exterior profile as the regular 3D-STED but with a nonzero-intensity center is achieved, which depletes the fluorescence signal at the center. In this way, we separate and detect the background fluorescence from the signal one. For clarification, we refer to the phase combination generating a regular 3D-STED image (3D-STED_on_; *R*) as “signal-on mode” and the phase combination generating a background 3D-STED image (3D-STED_off_; *B*) as “signal-off mode”, respectively. The 3D-STED_off_ (*B*) image can be subtracted from the 3D-STED_on_ (*R*) image with a subtraction factor *μ*, resulting in a background-free super-resolved 3D-STED image (DPS-STED; *S*), that is

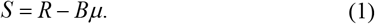

The subtraction factor *μ* corrects for the differences in background pixel values between the two sequentially recorded 3D-STED images. It is related to the different pixel dwell time and photobleaching effect in the experiments. The estimator *μ\** for *μ* is obtained by utilizing the least-squares method that minimizes 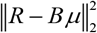. Specifically, *μ\** can be calculated by (Supporting Information, Section 8)

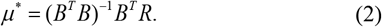

To demonstrate the background suppression capability of our method, we imaged fine subcellular structures in one and two colors within cells, for example, microtubules (Figure 2, Movie S1), mitochondria in U2OS cells (Figure 3, Movie S2), and the nuclear pore complex (NPC) and Golgi apparatus in Vero-B4 cells (Figure 4, Movie S3). The detailed imaging parameters are presented in Table S1. To allow nearly isotropic resolution, the power ratio between STED*_z_* and STED_*xy*_ beams was 4:1, enabling ~65 nm 3D resolution when the total STED laser power was ~210 mW (Figure S4). Note that a background 3D-STED frame was acquired immediately after recording a regular one at the same axial position using the same laser powers but with a shorter pixel dwell time to minimize the stage drift between the two frames. It also helps reduce the photodamage and the total imaging time. To compensate for the shot noise due to the shorter dwell time, we processed the background 3D-STED frames with a Gaussian filter of standard deviation *σ* = 11 pixels before subtraction.

**Figure 2.**
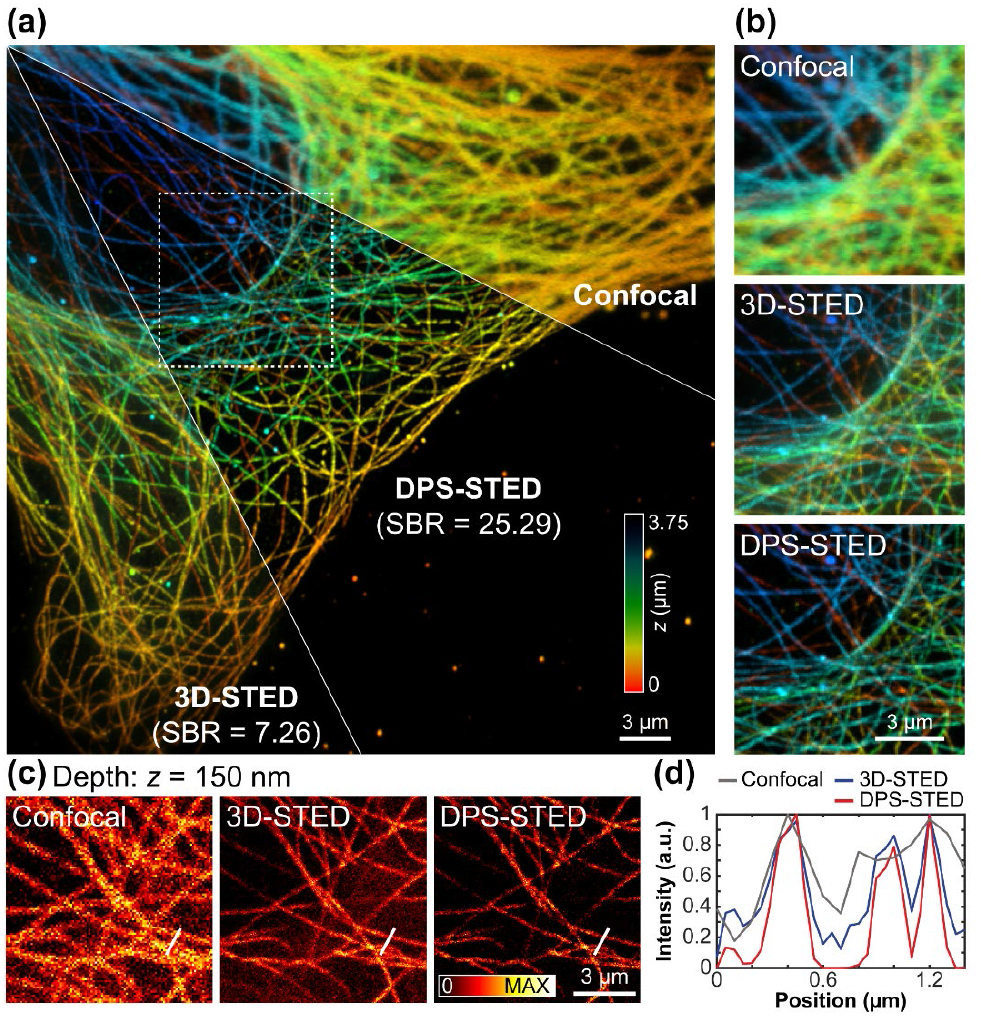
Images of microtubules in U2OS cells. (a) 3D distributions of tubulins (labeled with STAR ORANGE) are imaged under the confocal, regular 3D-STED, and DPS-STED configurations. The axial position of the microtubules is indicated using a rainbow colormap. (b) Magnified views from the region enclosed in the white dashed box in (a) under different configurations, (c) The *xy* views at *z* = 150 nm in (b). (d) Intensity profiles along the white solid lines in (c) show that DPS-STED reduces the background and neighboring filaments are better distinguishable.

**Figure 3.**
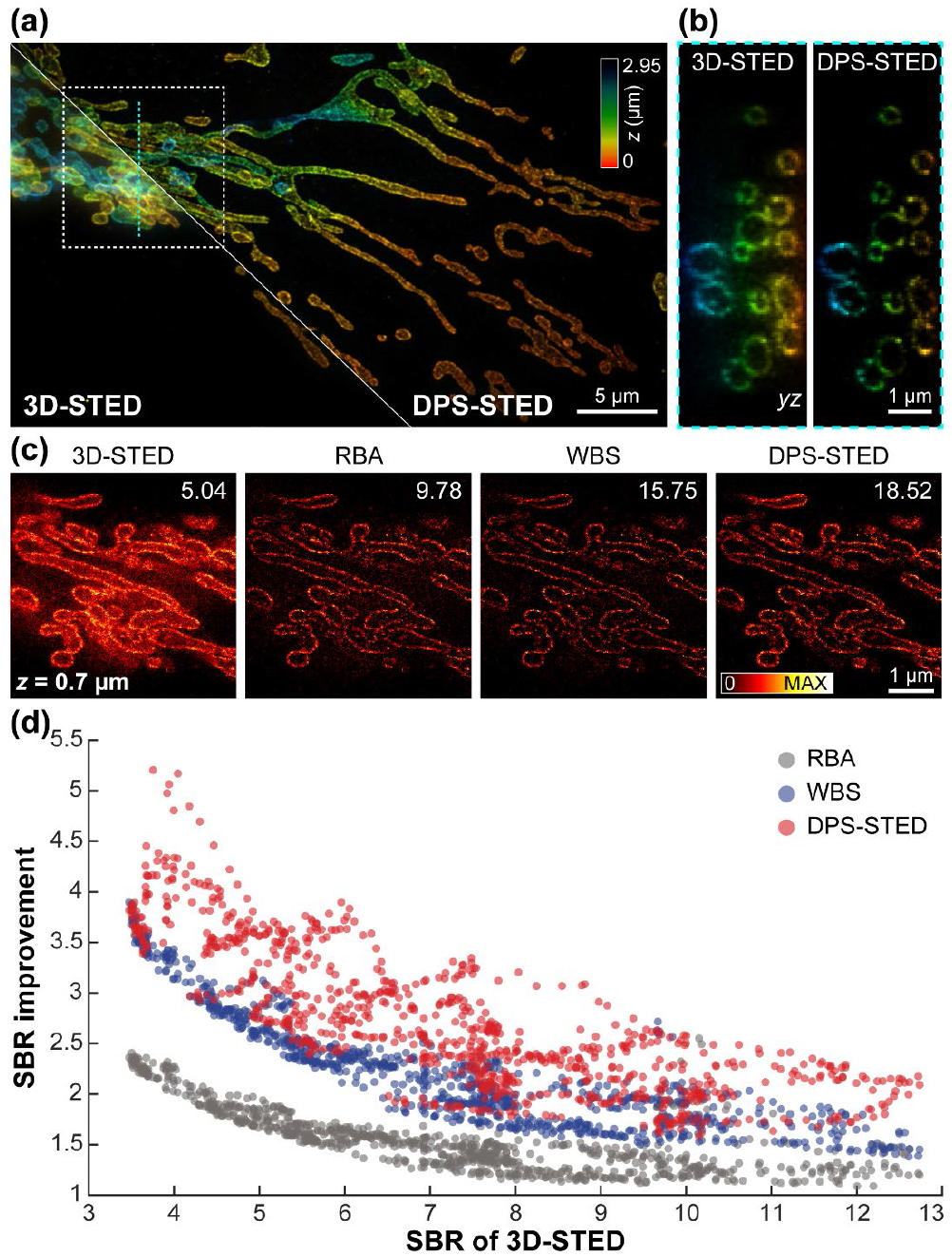
Images of mitochondria in U2OS cells. (a) 3D distributions of TOM20 (labeled with STAR RED) are imaged under the regular 3D-STED and DPS-STED configurations. The axial position of the mitochondria is indicated using a rainbow colormap. (b) The *yz* view of the cyan dashed line (120 nm wide) in (a) under different configurations. (c) The *xy* views from the region enclosed in the white dashed box in (a) at *z* = 0.7 μm under the regular 3D-STED, RBA (rolling ball radius: 3 pixels), WBS (full-width-half-maximum: 3 pixels), and DPS-STED. The SBR value is shown in the upper right corner of each *xy* view. (d) SBR improvement of different background subtraction methods. A total of 884 examples like (c) at different axial positions are used for analysis. See Figures S6 and S7 for details.

**Figure 4.**
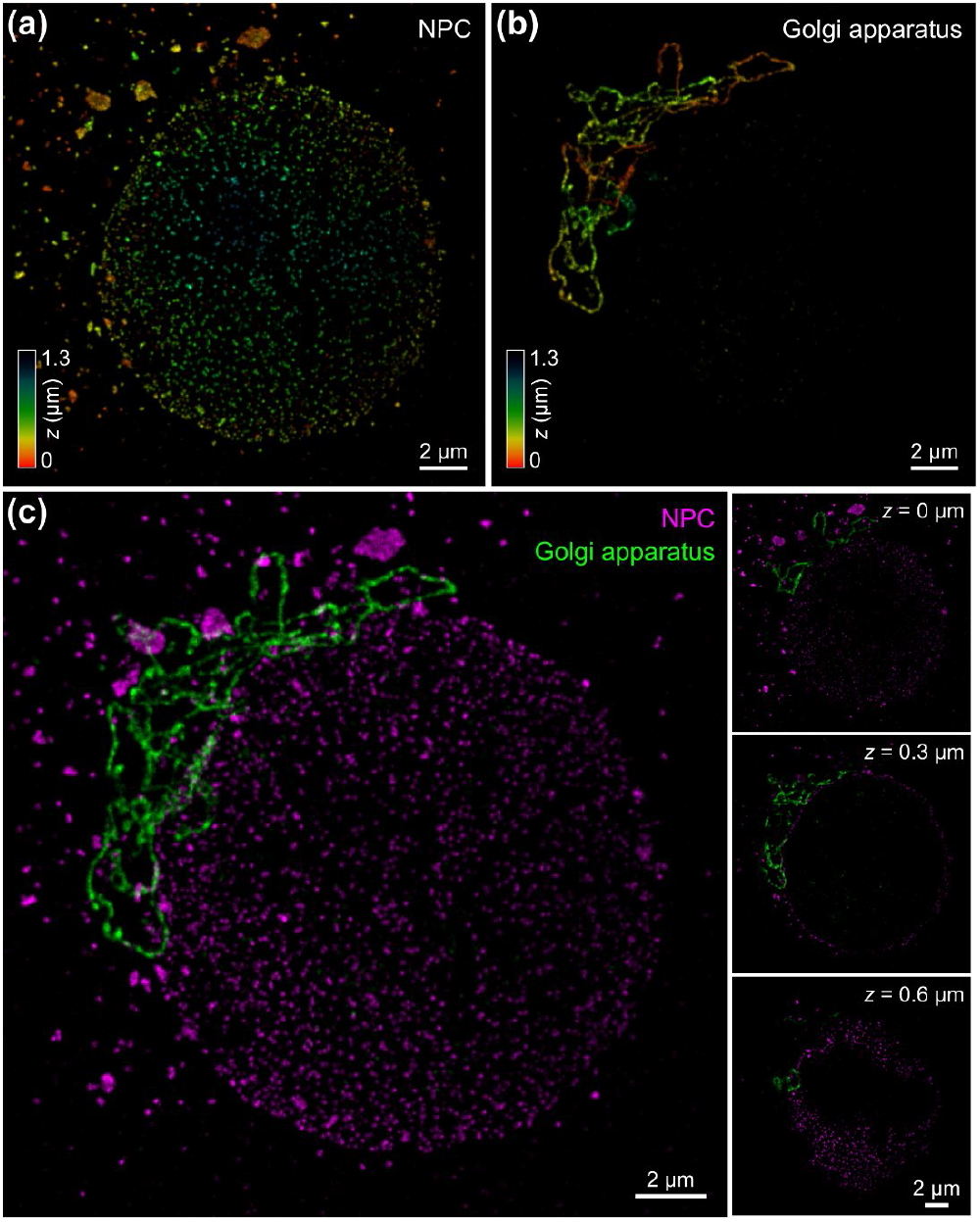
Dual-color images of NPC and Golgi apparatus in Vero-B4 cells using DPS-STED. (a) and (b) are 3D distributions of NPC (MAB414 labeled with STAR RED) and Golgi (Giantin labeled with STAR ORANGE). Rainbow colormap denotes axial position. (c) Merged image of two labels shown in (a) and (b). The *xy* views at three different axial positions are shown on the right.

In Figure 2a–b, 3D distributions of STAR ORANGE-labeled tubulins in U2OS cells are imaged under the confocal, regular 3D-STED, and DPS-STED configurations. In the DPS-STED image, the background signal is reduced dramatically, especially in the axially extended regions located near the nuclear envelope. The background suppression capability of DPS-STED is further revealed by plotting a line profile (Figure 2d) across the image (white solid lines in Figure 2c). The neighboring filaments are more clearly distinguished in DPS-STED. To quantitatively assess the background suppression performance, we calculated the SBR, marked in Figure 2a, of the two 3D-STED imaging modalities. Here, we defined SBR as the ratio of the averaged foreground signal to the averaged background. The mask used to delineate the foreground and background pixels was generated using the Otsu method^26^. Compared with regular 3D-STED (SBR = 7.26), DPS-STED (SBR = 25.29) achieved a >3-fold improvement.

We next imaged mitochondria in U2OS cells by immunolabeling TOM20 with STAR RED under regular 3D-STED and DPS-STED configurations (Figure 3a and Movie S2). The mitochondrial outer membrane in DPS-STED is more clearly distinguished in 3D due to the background suppression capability (Figure 3a-b). We further compared the performance of background reduction among DPS-STED, RBA, and WBS. The comparison results are shown in Figure 3c-d, which demonstrates that DPS-STED can obtain higher SBR than RBA and WBS in both low SBR and high SBR regions. Notably, as expected, the SBR improvement in the low SBR regions is more significant than that in the high SBR regions for all three methods.

In a more challenging demonstration, we performed dual-color imaging of NPC and Golgi apparatus in Vero-B4 cells using DPS-STED. In Figure 4a–b, the 3D distributions of NPC (MAB414 labeled with STAR RED) and Golgi apparatus (Giantin labeled with STAR ORANGE) are presented. As shown in Figure 4c, two spatially compact subcellular structures (NPC and Golgi apparatus) are super-resolved and reliably separated both laterally and axially with high SBR and high 3D resolution (see Movie S3).

Here we have demonstrated that DPS-STED exhibits improved SBR by imaging different subcellular structures in biological samples. DPS-STED nanoscopy is a differential imaging technique that accurately measures and removes background mainly arising from out-of-focus incomplete depletion in 3D-STED imaging experiments (Figure S5). To record the background, the key of DPS-STED is to switch the phase modulation dynamically by changing the handedness of the vortex phase mask and the inner radius of the top-hat phase mask. This modulation can directly be implemented via SLM. To guarantee that the recorded background contains only the background and no useful signal, it is of essence to modulate both the vortex and the top-hat phase masks. However, in an extreme case when the power of the STED*_xy_* beam is much higher than that of the STED_*z*_ beam (*e.g*, 2D-STED), changing the handedness of the vortex phase mask is sufficient, which brings more flexibility to the STED system in which fixed phase masks instead of an SLM is applied. Thus, we can introduce DPS-STED to various depletion power distributions between the vortex and the top-hat phase masks to meet the resolution requirements of a given imaging task. Besides, to accurately remove the background, we estimate the subtraction factor using the least-squares method by minimizing the norm of the background-subtracted image. As a result, compared with the psSTED method, DPS-STED enables that the pixel dwell time of the background image becomes much shorter than that of the regular 3D-STED image (Figure S8), thereby accelerating the imaging. Shorter pixel dwell time resulting in lower background intensity can be accommodated when using Gaussian filtering on the background image before subtraction to remove statistical noise. Furthermore, compared with software-based background subtraction algorithms (RBA and WBS), DPS-STED can obtain better background-free images with accurate background estimation.

To summarize, we have presented a novel method termed DPS-STED to suppress background in AO 3D-STED nanoscopy. Compared with other background suppression methods, through the combination of AO strategy, DPS-STED is easier to implement, more flexible, and more effective. Additionally, the DPS-STED method can be applied to 3D-STED live-cell imaging^25^ and fluorescence correlation spectroscopy (FCS) applications^18^ to improve SBR. Moreover, the subtraction factor determination method is also suitable for other differential background suppression methods for super-resolution microscopy^15, 16^.

## ASSOCIATED CONTENT

### Supporting Information

The Supporting Information is available free of charge at http://pubs.acs.org.

Detailed schematic of custom-built multicolor AO-3D-STED nanoscope setup; detailed process chart of the AO strategy used for STED imaging and effects of AO on the imaging performance; quantification of 3D resolution; background sources in 3D-STED nanoscopy; background reduction using RBA, WBS, and DPS-STED; imaging parameters for all images presented in the main text; cell culture and immunofluorescence staining; calculation of subtraction factor; and methods of image processing and visualization (PDF)
The confocal, regular 3D-STED, and DPS-STED imaging of microtubule in U2OS cells in Figure 2 (MP4)
The regular 3D-STED and DPS-STED imaging of mitochondria in U2OS cells in Figure 3 (MP4)
The dual-color DPS-STED imaging of NPC and Golgi apparatus in Vero-B4 cells in Figure 4 (MP4)

## AUTHOR INFORMATION

Complete contact information is available at: http://pubs.acs.org.

### Notes

The authors declare no competing financial interest.

## ACKNOWLEDGMENTS

This work was funded by the National Natural Science Foundation of China (92050115), Natural Science Foundation of Zhejiang Province (LZ21F050003), “Leading Goose” R&D Program of Zhejiang (2022C01077), and the Fundamental Research Funds for the Central Universities (226-2022-00137).

